# Size Matters: Metastatic cluster size and stromal recruitment in the establishment of successful prostate cancer to bone metastases

**DOI:** 10.1101/235911

**Authors:** Arturo Araujo, Leah M. Cook, Conor C. Lynch, David Basanta

## Abstract

Prostate cancer (PCa) impacts over 180,000 men every year in the US alone with 26,000 patients expected to succumb to the disease (cancer.gov). The primary cause of death is metastasis, with secondary lesions most commonly occurring in the skeleton. Prostate cancer to bone metastasis is an important yet poorly understood process that is difficult to explore with experimental techniques alone. To this end we have utilized a hybrid (discrete-continuum) cellular automata (HCA) model of normal bone matrix homeostasis that allowed us to investigate how metastatic PCa can disrupt the bone microenvironment. Our previously published results showed that PCa cells can recruit mesenchymal stem cells (MSCs) that give rise to bone building osteoblasts. MSCs are also thought to be complicit in the establishment of successful bone metastases (1). Here we have explored aspects of early metastatic colonization and shown that the size of PCa clusters needs to be within a specific range to become successfully established: sufficiently large to maximize success but not too large to risk failure through competition amongst cancer and stromal cells for scarce resources. Furthermore, we show that MSC recruitment can promote the establishment of a metastasis and compensate for relatively low numbers of PCa cells seeding the bone microenvironment. Combined, our results highlight the utility of computational models that capture the complex and dynamic dialogue between cells during the initiation of active metastases.

## 1. Introduction

Metastasis is a deadly process that necessitates cancer cell intravasation, circulation, extravasation and establishment in the new location (2). Prostate cancers typically metastasize to the skeleton where, upon establishment, they generate osteolytic and osteoblastic lesions that are painful and contribute greatly to the morbidity of the disease (3). In the bone microenvironment, prostate cancer cells manipulate adult bone forming osteoblasts (aOBs) and adult bone resorbing osteoclasts (aOCs) to yield growth factors and space for expansion and PCa growth in a process described as the “vicious cycle” (4). To be successful, PCa cells have to adapt to the bone environment and take advantage of the available resources in a Darwinian fashion (5). The key properties of these PCa cells include the recruitment of MSCs; pluripotent cells that can produce a population of precursor osteoclasts (pOBs). These key cells contribute to the vicious cycle by expressing receptor activator of nuclear kappa B ligand (RANKL) – a signaling molecule that promoted the activity of bone-resorbing osteoclasts. Increased bone resorption results in the release of bone derived factors that in turn drive PCa growth thus completing the cycle (6). The process by which prostate cancer to bone metastases successfully and establish in the bone microenvironment is less clear. We suggest that understanding and quantifying the dynamics of molecular and cellular interactions involved in the establishment of active metastases, i.e. not dormant disseminated tumor cells (DTCs) using computational modeling can provide new insights into this aspect of the disease with a potential for the development of interventional therapeutic strategies. For this purpose, we have integrated relevant biological parameters into a computational model that can address this clinically significant problem. Our model opens a window into the inner dynamics of the cells, their heterogeneity, their interactions and their behaviors across time and space; features that are difficult to be determine using traditional experimental approaches. The current paradigm of how cancer cells extravasate in the metastatic microenvironment and form an active lesion revolves around the interactions between cancer cells and the surrounding microenvironment. It is now known that circulating tumor cells can travel in clusters and that, compared to single circulating cells, their ability to metastasize is up to 50-fold higher (7,8). It is currently very difficult to have accurate control on the cluster size of PCa cells used to seed a metastasis experimentally. Further, *in vivo* experimental approaches often do not afford the resolution to identify subtle changes that occur at key time points in the early stages of metastatic seeding. Computational models like the one we propose here, can tackle these issues by offering a control over the discretization of the tumor initiating cells and MSC recruitment, while allowing for a full high-resolution record of the dynamics of each population and how they change in response to these discrete changes. We believe that our approach can tease out important interactions between the metastatic tumor and the surrounding microenvironment leading to potentially useful biological and clinical knowledge and even novel avenues of research.

## 2. Computational model and experimental setup

Previously we used a computational model to study prostate cancer (PCa) to bone metastases that was parameterized with empirical/published cellular and molecular values (9). We focus on the Bone Modelling Unit (BMU), a 3.0x2.0mm area where bone remodeling takes place. Figure 1 describes the model including the main cell types and signaling molecules and the rules that govern the behavior of the cells. Briefly we will describe some of the key elements of the model here. We considered 6 different cell types, including 5 residents of the BMU: Osteoblasts (OBs), Osteoclasts (OCs), precursor Osteoblasts (pOBs), precursor Osteoclasts (pOCs), MSCs, as well as prostate tumor cells (PCa), capable of recruiting MSCs and producing TGFβ. We also considered the interactions between the key cell types and the microenvironmental factors that control those interactions, which were defined as follows:

- Bone. Bone is the richest reservoir of TGFβ in the human body (700pg/mg of bone tissue). We have modeled bone explicitly as static cells that, when resorbed, disappear from the domain and release bone derived factors (BDF) and TGFβ.
- MSCs, pOBs and OBs. Bone generating osteoblasts (pOBs) are derived from MSCs. MSCs undergo asymmetrical division and pOBs proliferate in response to TGFβ, ultimately differentiating into bone matrix producing OBs, a process mediated by factors such as BMP-2. In our model, pOBs express RANKL, migrate towards and expand clonally in response to TGFβ, and finally differentiate into OBs after 14 days. As adult OBs, the modeled cells seek TGFβ and bone. If in contact with bone, they lay down bone matrix with an active lifespan of 75 days.
- pOCs and OCs. pOCs are derived from myeloid precursors and, in response to RANKL, undergo cellular fusion to generate mature OCs. OCs resorb the bone matrix leading to the release of BDFs and TGFβ. We have explicitly modeled these processes. pOCs are recruited by RANKL from the vasculature, and have a lifespan of 2 days. Once on the bone surface, they will fuse together provided that the local levels of RANKL are high while those of TGFβ are low. A minimum of 3 pOCs (usually 5 or more) can fuse to form an OC. OCs have a lifespan of 14 days, in which their singular function is to resorb bone thereby releasing TGFβ and BDFs. Based on the amount of TGFβ present in bone, we have calculated that a single osteoclast measuring 100μm in diameter will resorb approx. 10μm^3^of bone per day. Given the density of bone as 1500mg/Kg, we estimate that an osteoclast can resorb 1.17x10^−3^mg/day. With the concentration of TGFβ in bone being 5ng/mg, we calculated that a mature osteoclast can generate up to 0.00558 ng of TGFβ per day.
- PCa. Based on our empirical as well as published data, we engineered the PCa cells to express TGFβ ligands and receptors. Importantly, the level of PCa TGFβ (5x10^−12^pg/day) is approximately 1000-fold less than that generated by bone resorbing OCs (5x10^−9^pg/day). This ensures the reliance of the prostate metastases on TGFβ released from the bone. In the computational model, we have described the TGFβ producing PCa metastases as agents chemoattracted to the BMU, with an ability to recruit MSCs, based on our empirical studies. PCa replication potential is proportional to the availability of bone-derived factors and, if not bathed with these essential factors, PCa cells will die within 14 days. The PCa metastases are the only ones that can destroy the canopy of the BMU as they grow. We have considered that PCa promote OB differentiation, a phenomenon that is not noted in lytic lesions.
- The microenvironment. We model the BMU as a 300x200 pixel area, where the sub-indices i and j denote the position of the cells in the grid that produce or interact with key molecules. The key molecules TGFβ, RANKL and BDFs are generated by the behaviors and interactions amongst the cellular components; and are characterized by partial differential equations that are subsequently discretized and applied to a grid. TGFβ is produced by bone destruction (*α_β_B_i,j_*) and cancer cells (*α_c_C_i,j_*) in proportion to the local TGFβ concentration, with natural decay of the ligand (*σ_β_T_β_*); ensuring the density never exceeds a saturation level, m_0_. TGFβ has pleiotropic effects on osteoblasts, osteoclasts and metastatic prostate cancer cells. Low concentrations of TGFβ stimulate osteoclastogenesis but high concentrations inhibit the process; illustrating the biphasic effects of this growth factor even on the same cell type. TGFβ is governed by the following differential equation:

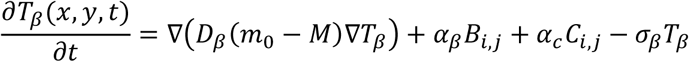
- RANKL *R_L_* is produced by precursor Osteoblasts, *α_L_O_i,j_*, in proportion to the local RANKL concentration, with natural decay of the ligand, *σ_L_R_L_*; ensuring the density never exceeds the saturation level n_0_. The concentration of RANKL is determined by:

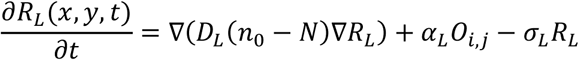
- Factors *F_B_* are released by bone destruction, *α_B_B_i,j_*, in proportion to the local factor concentration, with natural decay of the factors, *σ_B_F_B_*; ensuring the density never exceeds the saturation level p0. As such, the dynamics of the bone related factors are calculated through:

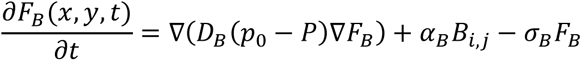

**Figure 1.**
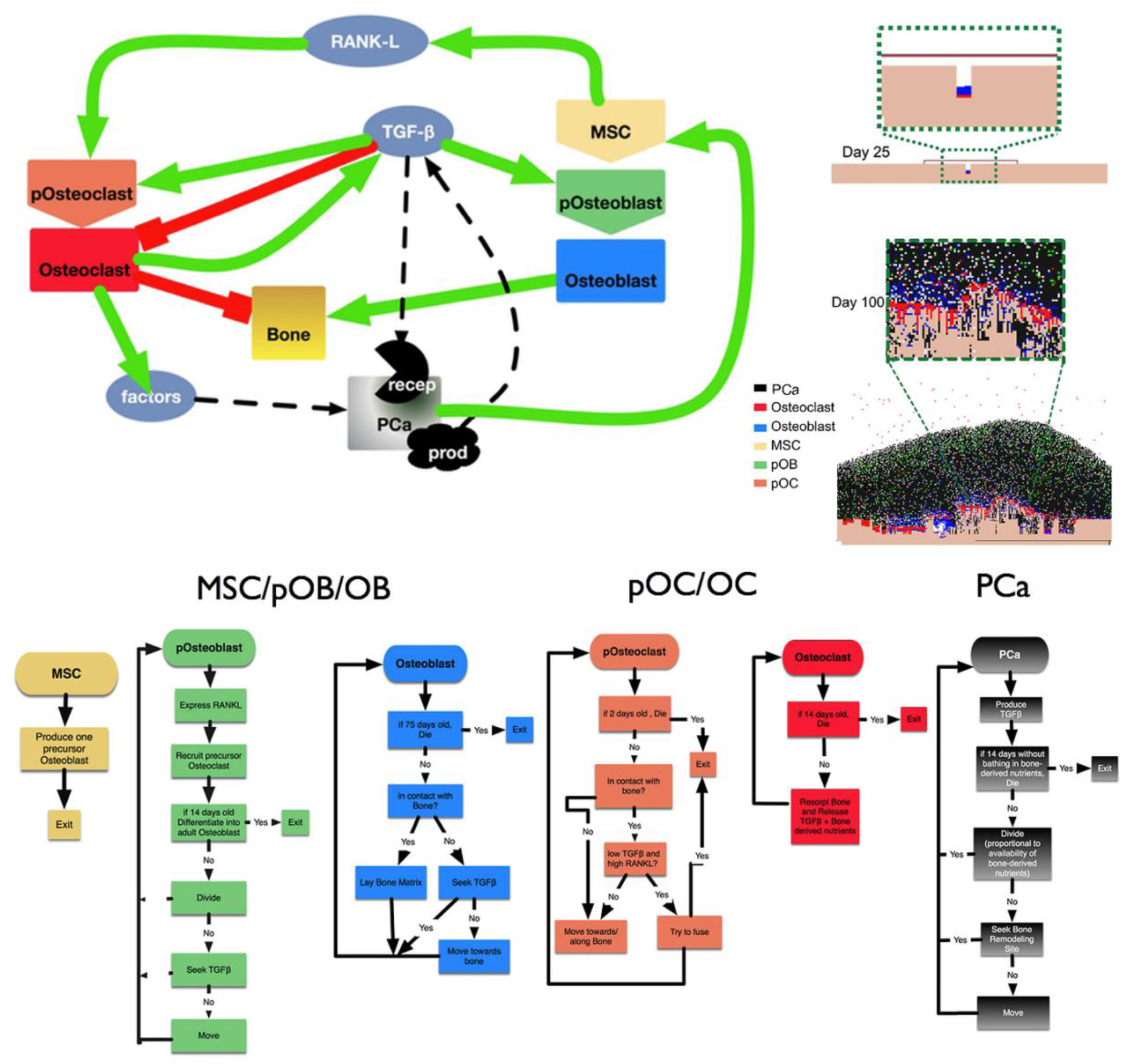
Top left panel: Based on our previous model (9) we considered 7 different cell types: resident and active cells of the bone stroma (mesenchymal stromal cells (MSCs), precursor osteoblasts (pOB), adult osteoblasts (aOB), precursor osteoclasts (pOC), mature osteoclasts (aOC), bone cells) and prostate cancer cells. Homoeostasis in the bone microenvironment is regulated by the signaling factors TGFβ, RANKL and bone derived nutrients. These molecules are generated by the behaviors and interactions across the cellular players; and are characterized by partial differential equations that are subsequently discretized and applied to a (300x200 pixel) grid. Top right panel. A representative bone metastatic prostate cancer vicious cycle simulation at day 25 and 100. Bottom panel. The rule sets guiding the behavior of each of the cell populations used in the HCA model. Adapted from Araujo et al., (9).

Periodic boundary conditions were considered only for the left and right sides of the microenvironment, while no-flux boundaries were imposed on the top and bottom of the 2D grid. This HCA model recapitulates bone remodeling and homeostasis and, when a tumor cells is seeded, it can potentially lead to the establishment of a vicious cycle (figure 1 top right) (9). We previously explored and biologically validated several hypothesis regarding treatment of metastatic prostate cancer with this HCA model (9,10). Here we turn to the earlier stages of metastasis and have 1) varied the initial seeding of PCa cells from 1 to 10, 100, 250, 500, and 750 to examine the impact of PCa cluster size on successful metastasis initiation and 2) altered the rate of MSC recruitment (1 to 1/1000 in 10-fold reductions) to determine whether MSCs play a role in establishment of PCa bone metastases. We have also investigated the effect of the combination of both variations. In the computational model, each cell is represented individually as an agent (11) (12) and realistic biological behavior emerges from the interactions of the cells and the microenvironment (9). The system recapitulates the normal bone modeling process and the dynamic balance between bone regenerating MSCs, precursor osteoblasts (pOBs), adult osteoblasts (aOBs) and the bone resorbing precursor osteoclasts (pOCs) and adult osteoclasts (OCs) (Figure 1). Factors such as transforming growth factor β (TGFβ), RANKL and bone derived factors stored in the bone matrix tightly regulate the process of bone remodeling and homeostasis (Figure 1). In the computational model, each cell type responds to TGFβ levels in an either directly proportional (1+*Log*(*TGFβ*)) or inversely proportional (-1*Log*(*TGFβ*)) manner. Computational models were seeded with homogeneous prostate cancer cells that expressed the TGFβ receptor and ligand (TRP). To provide evidence in support of a key hypothesis, simulations were also performed with prostate cancer cells that express the ligand alone (TP). For the HCA model, we consider cellular intrinsic behaviors and the impact of TGFβ on these behaviors. Empirical, experimental and theoretical parameters were used to fuel equations (Table 1). The rate of pOB division is inversely affected by TGFβ and we assume that the effect is logarithmic. By the same token, the fusion rate of pOC is also affected inversely proportional to the availability of TGFβ. Tis is implemented by through equation:

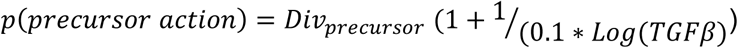

where 0< TGFβ <1 and *Div_precursor_* is substituted in the case of precursor osteoblasts, *Div_p0B_* is the maximum rate of pOB division, and in the case of precursor osteoclasts, *Div_p0C_*, the maximum rate of pOC fusion in the absence of TGFβ. Furthermore, once fused, the probability of aOC survival also depends proportionally on TGFβ, as described by:

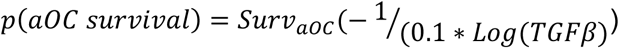

where 0< TGFβ <1 and *Surv_a0C_* is the maximum percentage of survival for aOCs. Finally, TRP and TP division in response to bone derived factors was modeled as follows:

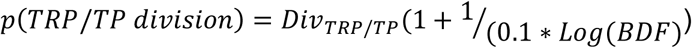

**Table 1.**
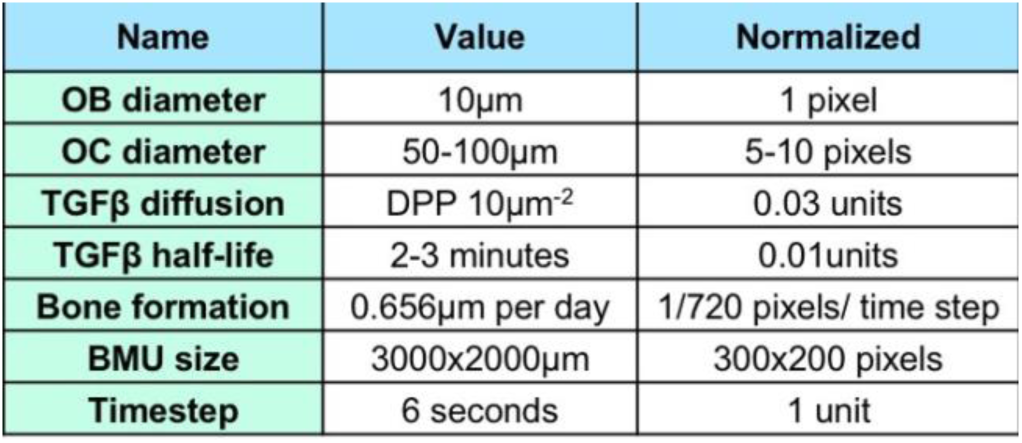
Select parameters used to generate the HCA computational model and adapted from (9).

Based on empirical data obtained with TRP cells such as the prostate cancer cell line, PAIII (9), we assume that TRP cells have a maximum division rate of once every 1.5 days and a lifespan without contact with Bone derived Factors of 14 days. We assume that TP has a maximum division rate of once 1.75 days, and a lifespan without nutrients of 10 days, calculated as having a cost for producing TGFβ ligand but not benefiting from TGFβ contained within the bone derived nutrients.

PCa cells have the potential of disrupting bone remodeling. The secretion of the TGFβ ligand by PCa cells, albeit at lower concentrations than those that can be found in the bone, can result in the recruitment of MSCs followed by unregulated pOB expansion that express RANKL, thus causing an influx of osteoclast precursors. This in turn leads to increased osteoclast formation and bone destruction. Low concentrations of TGFβ stimulate osteoclastogenesis but high concentrations inhibit the process; illustrating the biphasic effects of this growth factor even on the same cell type (13). Our group has shown that the release of bone-derived nutrients and sequestered growth factors from the bone matrix, such as TGFβ, promotes the survival and growth of the metastatic prostate cancer cells, thus perpetuating the vicious cycle (9).

## Results and Discussion

### Metastatic PCa clusters (>100 cells) establish less frequently in the bone microenvironment

We first investigated the question of whether the number of disseminated prostate cancer cells has an impact on the establishment of a successful metastasis. A single proliferating PCa cell can generate potentially generate a metastatic lesion, later branching into a more heterogeneous tumor through microevolution (14). Theoretically, it is posited that larger PCa clusters would be capable of more efficiently establishing bone metastases (15). To determine the impact of PCa cluster size on metastatic success, we designed a series of experiments to systematically vary the initial seeding of TGFβ-responsive PCa cells from 1 to 750. Each experiment consists of a set of 50 simulations. For statistics on the cell populations we only considered those simulations that successfully established a metastasis by day 125. Our data was assessed at days 125 and 250 since we previously determined that these points are relevant in a biologically scaled mouse model of bone metastatic prostate cancer (10).

The average response of each microenvironmental cell at day 125 (blue) and at day 250 (red) to an initial seeding of 1, 10, 100, 250, 500 and 750 PCa cells, illustrated in Figure 2, as circle whose diameter represents the standard deviation. Also shown in the right upper corner of each scenario the number of successful metastases out of a total of 50 simulations. These results show that the success of metastasis is increased up to a threshold of approximately 100 PCa cells after which our results show a decrease in the likelihood of successful metastatic initiation as we increase the size of the initial PCa cluster. The size of the tumor burden correlates with the initial seeding up to 100 cells. Further, the seeding number has a direct effect on the microenvironmental cells, increasing the magnitude of the vicious cycle via a rapid expansion of pOBs and aOCs. Interestingly, we observed that at day 125, PCa clusters of between 10 and 100 impacted bone volume significantly. Increasing the PCa cluster size greater than 100 cells had no further impact on PCa burden or PCa induced bone disease. By day 250, the PCa burden and microenvironmental responses were similar in all successful metastasis simulations.

**Figure 2.**
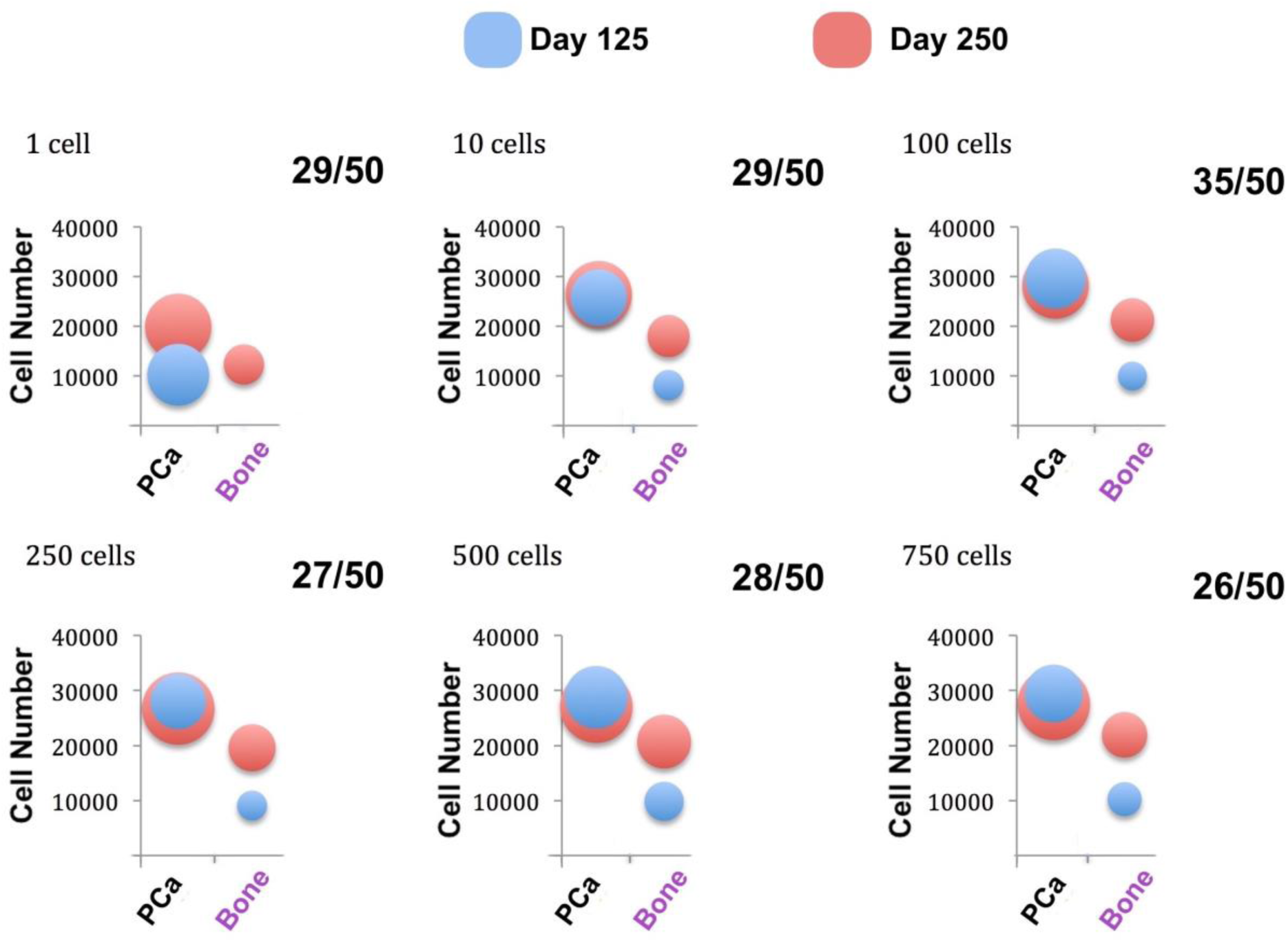
Microenvironmental response at day 125 and at day 250 to an initial seeding of 1, 10, 100, 250, 500 and 750 PCa cells. The success of metastasis (top left of ratio in each experiment) increases up to a threshold at which the number of cells in the metastatic bundle does not translate into higher chances of metastatic success. A similar assessment can be made of the importance of that metastatic bundle size on the final tumor size.

### MSC recruitment can compensate for other deficiencies in the PCa phenotype

Evidence suggests that that MSCs significantly contribute to prostate cancer induced osteogenesis (9), but whether MSCs impact the establishment of prostate bone metastases remains unknown. To address this with our computational model, we varied the MSC content in the bone microenvironment (1 to 1/1000 in 10-fold reductions) resulting from the recruitment by the overexpression of TGFβ by tumor cells. Figure 3 describes the response of the microenvironment at day 125 (blue) and at day 250 (red) to the reduction of MSC recruitment of 1/10, 1/100 and 1/1000 (horizontally); how MSCs vary depending on the initial seeding of PCa cells of 10, 100 and 1000 (vertically); and the combined effect of both. Results from Figure 3 show that the inhibition of MSC recruitment reduces tumor burden in simulations where lower numbers of PCa cells are seeded. However, increasing the PCa cluster size can overcome the effects of lower rates of MSC recruitment. As expected, reducing the number of MSCs recruited to the cancer-bone microenvironment also reduced the numbers of osteoblasts, osteoclasts and bone volume regardless of variations in PCa cluster size.

**Figure 3.**
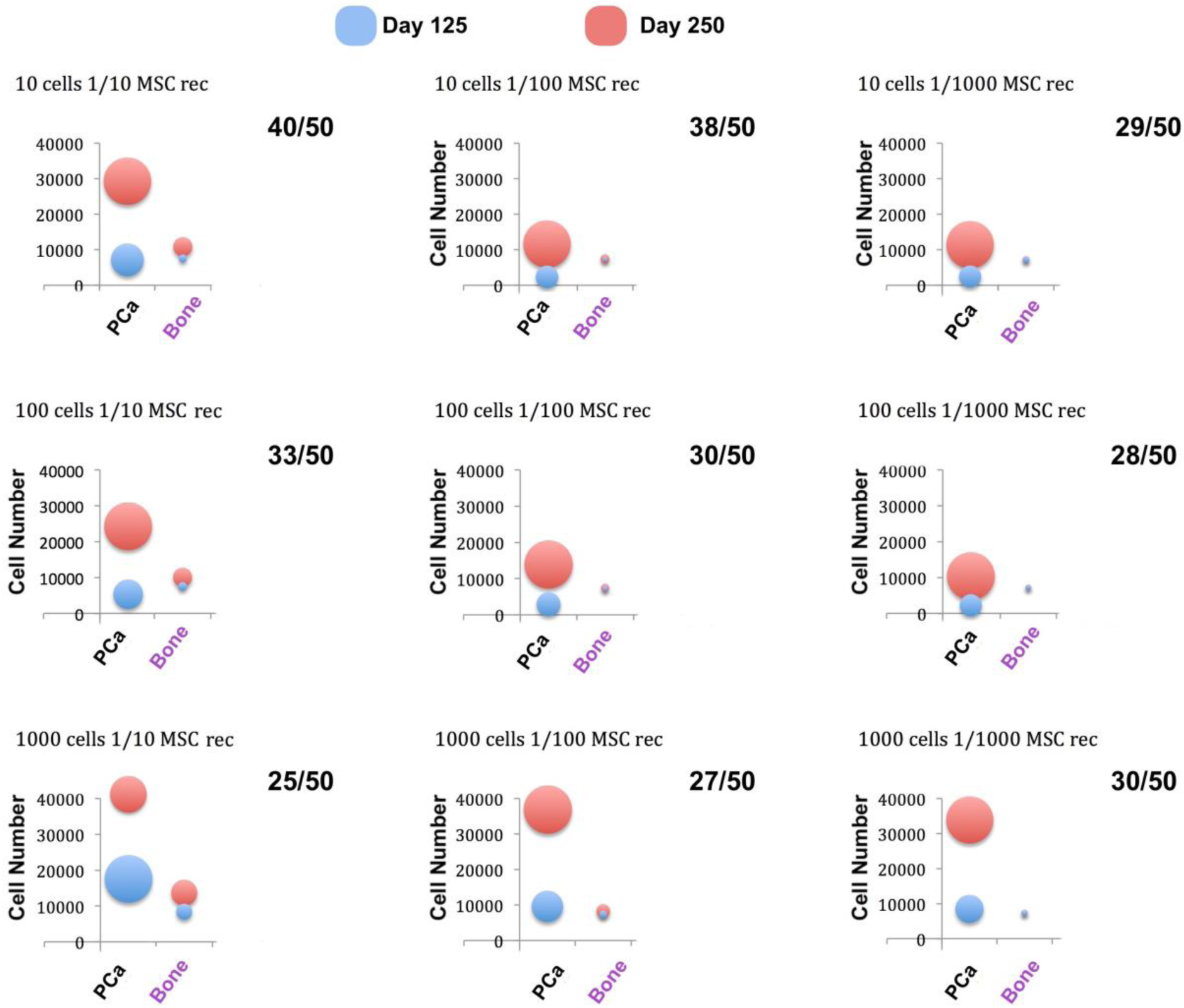
Microenvironmental response at day 125 and at day 250 to the reduction of MSC recruitment of 1/10, 1/100 and 1/1000 (horizontally); and the initial seeding of PCa cells of 10, 100 and 1000 (vertically). The inhibition of MSC recruitment reduces tumor burden in low seedings. However, as the number of PCa seeding cells is increased, tumor is also exacerbated.

### Competition for resources is critical for the successful establishment of PCa metastases

Our results show that in some cases, higher PCa cluster sizes seeded into the bone microenvironment counter-intuitively failed to successfully establish metastases. In these scenarios, we posit that the PCa cells are competing for TGFβ with other TGFβ regulated microenvironmental cells. If so, this would mean that if the PCa cell number are initially increased, then the proportion of OCs and OBs would be reduced since there are less bioavailable nutrients; thus, countering the added benefit that a large seeding could have on initiating the vicious cycle. To test this hypothesis, we can dissect the inner dynamics of a representative simulation, and compare it to the same simulation, replacing PCa cells with non-consuming TGFβ-producing PCa cells; an example of a truly unique ability of computational simulation to isolate the response to a key change in a single variable, while keeping other factors constant (16).

Figure 4 shows the result of a representative control simulation of a TGFβ receptor expressing PCa 10-cell cluster seeding (TRP) with normal MSC recruitment rates (left); and the same simulation seeded with a non-TGFβ consuming (TP) 10-cell PCa cluster (right). Although the cell populations look similar on the surface, there are subtle but important temporal differences in the cancer and stromal cell populations. The microenvironmental changes in the TRP PCa metastases are smoother, and osteocalstogenesis reaches a maximum, after which the tumor and the microenvoronmental cells collapse due to competition for TGFβ and nutrients. In the case of the TP PCa metastases, because there is an abundance of TGFβ and the tumor is not in direct competition with the microenvironment cells, the dynamics are more regular and their cycles more pronounced, suggesting more rapid but stable changes in the microenvironmental cells. Specifically, precursor osteoclasts and osteoblasts have a more pronounced phasic behavior, suggesting stability and the population of adult osteoclast (bottom) is reduced because excess TGFβ inhibits pOC fusion, in line with our theory. Although this would seem more advantageous for a tumor, however, we noted that only 15/50 simulations seeded with TP cells successfully metastasized, in contrast with the 29/50 successful TRP metastasis under the same randomness conditions. This suggests that although TRP cells are important for the intitation of a vicious cycle, the tumor might then benefit from evolving to a TP phenotype to sustain it.

**Figure 4.**
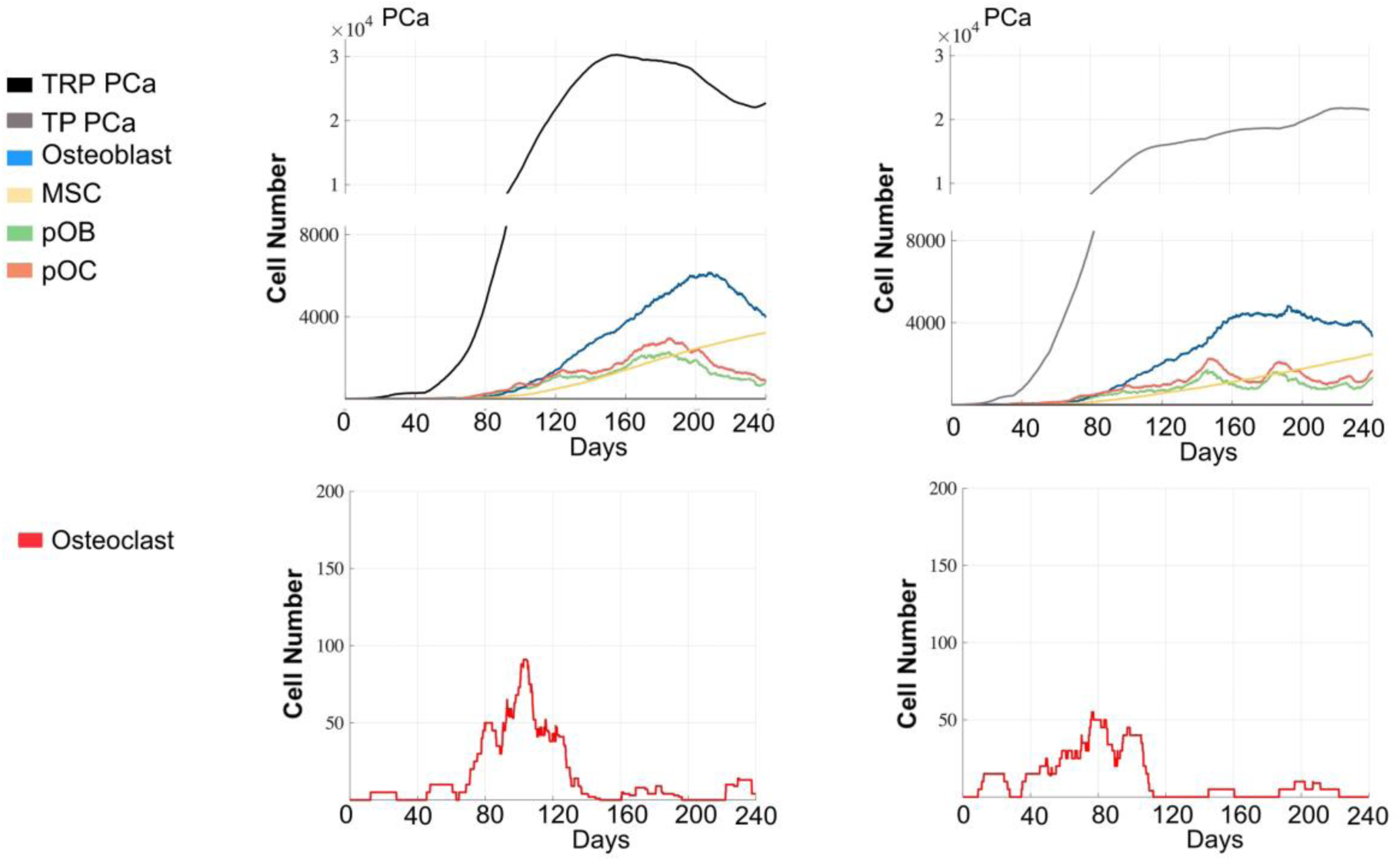
Comparing our previously described PCa cells (10 cell seeding) vs a different PCa cell line that is less sensitive to TGFβ (10 cell seeding). While the dynamics look similar on the surface, but there are subtle differences. The microenvironmental changes in the regular PCa case are smoother, and osteocalstogenesis is constant. In the less TGFβ sensitive case the dynamics are more irregular, suggesting dynamic changes in the microenvironmental cell populations. Only 15/50 simulations of TP were successful compared to the 29/50 of TRP.

Further, in our investigation of the role of MSC recruitment, we found that as the number of PCa seeding cells is increased, PCa tumor burden is reduced; countering the benefits gained by the reduction in MSC recruitment. This apparent tumor growth recovery could be explained by the absence of nutrient-consuming stromal cells, over-saturating the compartment with growth factors, helping PCa cells survive. To investigate this explanation, we can again explore the individual dynamics of representative simulations at the extremes of Figure 3: PCa 10 cell cluster seeding with normal MSC recruitment vs. PCa 1000 cell cluster seeding w. 1/1000 reduction in MSC recruitment rates (Figure 5). Being able to analyze the complete dynamics is key to teasing out interactions of significance that give rise to these complex behaviors. In this case, we discovered that, although these two simulations present similar numbers at days 125 and 250, the growth dynamics are quite different. In line with our explanation, results from Figure 5 suggest that the inhibition of MSC recruitment does reduce the microenvironmental burden, delaying the vicious cycle and while initially decreasing the tumor magnitude compared to baseline, the bio-availability of nutrients is enough to support PCa survival. Figure 5 demonstrated a slow progression towards an established vicious cycle in the 10 cell PCa cluster seeding with 1/10 MSC rate of recruitment. Compared to the simulations seeded by 1000 tumor cells and 1/1000 MSC recruitment rate scenario, we can observe that, in the latter case, the sheer number of PCa cells are able to initiate a vicious cycle on several occasions, but are unable to sustain it. The magnitude of the microenvironmental cells are reduced to very low levels, but the bio-availability of nutrients are sufficient to sustain the tumor for at least the duration of the simulation. This shows that the tumor growth has not a recovered, but can survive in contrast to high seeding of PCa cells and no manipulation of MSC recruitment. Importantly, the populations of microenvironmental cells as well as the bone burden are reduced by the inhibition MSC recruitment; regardless of the increase in PCA seeding compared to baseline.

**Figure 5.**
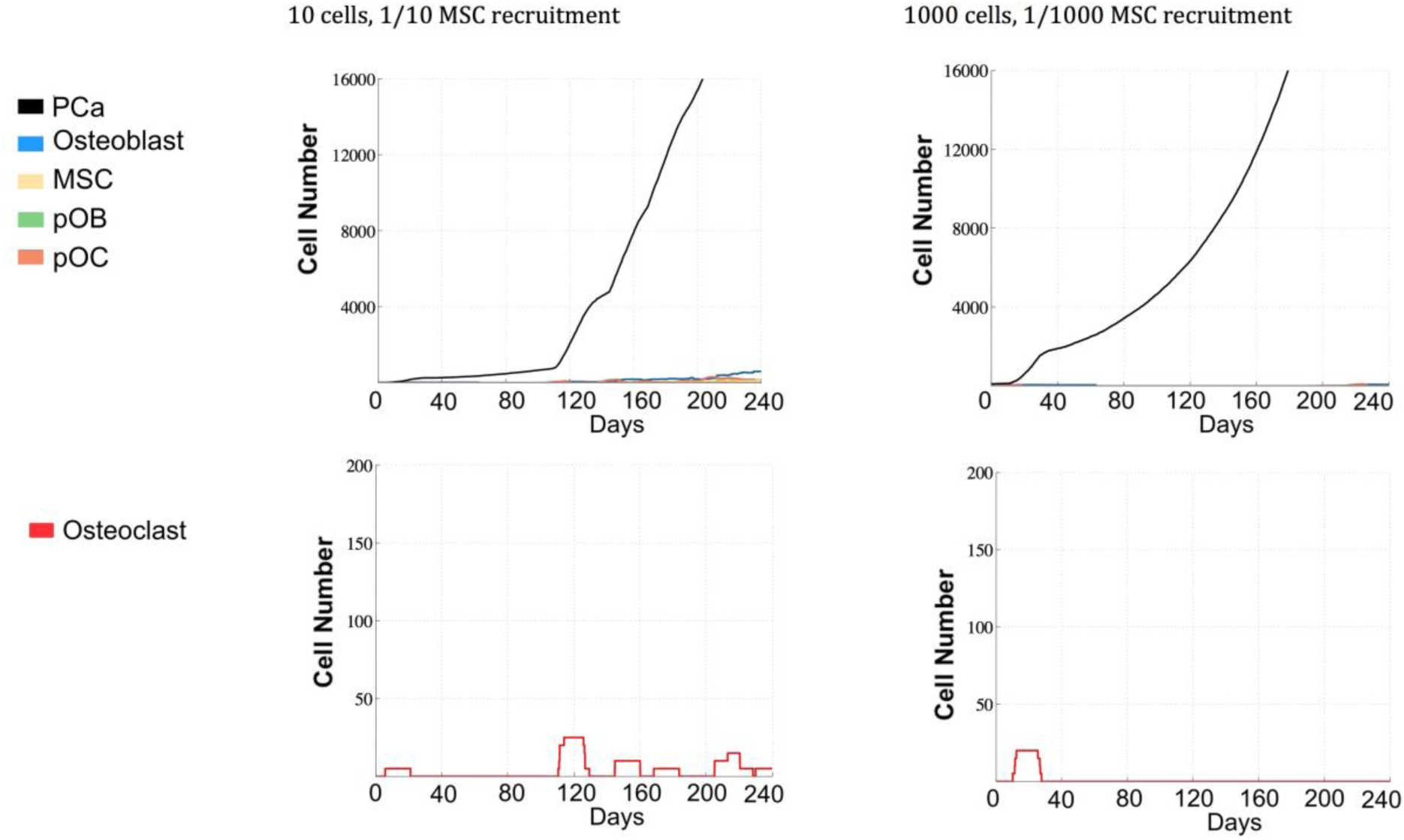
10 PCa cell seeding with normal MSC recruitment vs 1000 PCa cell seeding w. 1/1000 MSC recruitment. Although these two simulations present similar numbers at days 125 and 250, the inner dynamics are quite different. The inhibition of MSC recruitment reduces the microenvironmental burden, delaying the vicious cycle and initially decreasing the tumor magnitude.

## Conclusion

Herein, we have described an agent-based HCA model approach to address questions pertaining to the impact of PCA cluster seeding size in the establishment of successful metastases and whether bone stromal cells namely MSCs, impact the success of metastatic outgrowth. At this juncture, we have not considered questions pertaining to metastatic prostate cancer cells entering dormancy. Our results indicate that PCa cluster sizes up greater than 100 cells often fail to successfully establish metastases and that MSC recruitment to the site of the cancer seeding also contributes to the initiation of the vicious cycle. Based on our results we predict that PCa clusters greater than 100 cells fail to form metastases on occasion due to out competing cells of the local bone microenvironment for resources. Taken together, our results underscore the utility of computational modeling in tackling questions that are difficult to address with traditional *in vivo* approaches and providing outputs that can guide biological experimentation. Our data also demonstrates the important roles MSCs play in prostate cancer establishment and growth and indicate that inhibition of MSC recruitment/activity may be a viable therapeutic strategy for the treatment of bone metastatic prostate cancer.

## Acknowledgements

We would like to acknowledge Dr. Anderson from Moffitt’s Integrated Mathematical Oncology department for helpful discussions. AA, LC, CCL, and DB were partly funded by an NCI U01 (NCI) U01CA202958-01 and a Moffitt Team Science Award. AA was partly funded by a Department of Defense Prostate Cancer Research Program (W81XWH-15-1-0184) fellowship. LC was partly funded by a postdoctoral fellowship (PF-13-175-01-CSM) from the American Cancer Society.

